# Computer vision analysis of mother-infant interaction identified efficient pup retrieval in V1b vasopressin receptor knockout mice

**DOI:** 10.1101/2024.02.16.580409

**Authors:** Chortip Sajjaviriya, Fujianti, Morio Azuma, Hiroyoshi Tsuchiya, Taka-aki Koshimizu

## Abstract

Close contact between lactating rodent mothers and their infants is essential for effective nursing. Whether the mother’s effort to retrieve the infants to their nest requires the vasopressin-V1b vasopressin receptor axis has not been fully defined. To address this question, V1b receptor knockout (V1bKO) and control mice were analyzed in pup retrieval test. Because an exploring mother in a new test cage randomly accessed to multiple infants in changing backgrounds over time, a computer vision-based deep learning analysis was applied to continuously calculate the distances between the mother and the infants as a parameter of their relationship. In an open-field, a virgin female V1bKO mice entered fewer times into the center area and moved shorter distances than wild-type (WT). While this behavioral pattern persisted in V1bKO mother, the pup retrieval test demonstrated that total distances between a V1bKO mother and infants came closer in a shorter time than with a WT mother. Moreover, in the medial preoptic area, parts of the V1b receptor transcripts were detected in galanin- and *c-fos*-positive neurons following maternal stimulation by infants. This research highlights the effectiveness of deep learning analysis in evaluating the mother-infant relationship and the critical role of V1b receptor in pup retrieval during the early lactation phase.

**Highlights:**

(1) Mother-infant relationship was calculated as distances by computer vision and object detection analysis.
(2) V1b knockout female mice tended to avoid entering centerfield in a new cage compared to control female.
(3) Lack of V1b vasopressin receptor in mice facilitated the pup retrieval in early infancy, even with shorter distances of total movements.

## 1. Introduction

Feeding and protection of infants require a sustaining effort of the parents. Although maternal behaviors are pervasive across various animal species^1^, how the maternal behavior is initiated after parturition and maintained for a longer period is not yet fully understood. Mother-child interactions affect children’s survival in infancy and their physical and emotional development later in life ^2, 3^. Therefore, the mother’s health status, including mental health, is a key determinant of children’s wellbeing. In rodents, the maternal behaviors are comprised of a set of responses, such as nest building, crouching, licking/grooming, and retrieval of pups to the nest ^4^. Brain neural circuitry for these maternal behaviors involves two key areas in the hypothalamus, the medial preoptic area (MPOA) and its adjacent bed nucleus of the stria terminalis (BNST)^5, 6^. Previous studies have shown that galanin-containing neurons in MPOA are the central hub of maternal behavior in the early postpartum period^7–10^.

When stimulated by infants, MPOA neurons respond to increased estrogen, prolactin, and oxytocin (OT)^11–13^. Subsequently, the ventral tegmental area–dopamine pathway is activated ^4, 14^. Infant stimuli also activate the nucleus accumbens and the ventral pallidum, located laterally to the MPOA. Although OT receptor system play a crucial role in nurturing responses to pups^15^, a structurally related cyclic nonapeptide, arginine vasopressin (AVP), also mediates maternal care mainly through activation of central V1a receptors ^16, 17^. Applying the antisense of V1a receptor to the MPOA reduced pup retrieval, while the use of a V1a receptor antagonist to the BNST led to a decrease in defensive aggression in rat dams^18–20^. AVP is synthesized within the neurons of the paraventricular nucleus, supraoptic nucleus and BNST in the hypothalamus, and is released from the posterior pituitary to peripheral organs and broad areas in the central nervous system^21–23^. Among the three vasopressin receptor subtypes (V1a, V1b, and V2), the V1a receptor in vascular smooth muscle cells causes vasoconstriction and proliferation of vascular smooth muscle cells^24, 25^. In contrast, the V1b receptor is found in the anterior pituitary and central nervous system and is involved in hypothalamic-pituitary-adrenal axis activities^26, 27^. Both V1a and V1b receptors couple with heterotrimeric Gq protein and generating intracellular calcium signaling^28^. Bayerl et al. discovered that local infusion of a V1b receptor antagonist into the MPOA in rats resulted in augmentation of pup retrieval in a novel environment ^29^. How V1a and V1b receptors play different roles, despite of their coupling to the same Ca^2+^ pathway, in maternal behavior is not well understood.

Machine learning and computer vision enable the unbiased extraction of object features from images, facilitating the analysis of diverse behaviors in both experimental animals and humans ^30, 31^. By combining computer vision with deep learning, objects and their XY coordinates can be accurately identified against changing and challenging backgrounds. Furthermore, by applying deep learning, a vast amount of image and video data from animal experiments can be analyzed extensively beyond the capacity achievable by human efforts. Recently, animal behaviors, such as eating, drinking, and grooming, were successfully detected using computer vision and deep learning in mice and other animals ^32–38^. The main difficulty of analyzing mother-infant interaction lies in the ever-changing features of infants during the early phase of their development. During the first few days after birth, infants undergo noticeable daily changes in their skin color and size. Moreover, during pup retrieval, repeated and continuous accesses of a moving mother to infants makes it difficult to apply simple calculation of distances between the mother and multiple infants.

Here, we trained our deep learning model to localize mothers and their infants on the second and fourth days after parturition. Our established model was useful to detect exploratory behaviors of adult mice of both sexes in an open-field with and without infants. By analyzing V1bKO and WT mothers and their infants, the role of vasopressin in maternal motivation towards pup stimuli was studied during the early lactation period.

## 2. Materials and Methods

### 2.1 Animals

Mice were maintained in the animal room at 23 ± 1°C, 55 ± 10% relative humidity and a 12h light-dark cycle. Food and water were provided *ad libitum*. Generation of mice globally deficient in the V1b receptor was described previously ^26^. V1bKO and WT mice were maintained on a hybrid 129/Sv and C57BL/6 background. WT or V1bKO female mice at 12– 20 weeks of age were mated with male mice and kept in a clear polycarbonate resin (PC) cage (22 × 32 × 14 cm). Males were separated from the females prior to labor. Dams and pups were kept in the home cage throughout the study duration, except for the pup retrieval test. The cage bedding was kept for 10 days following labor to avoid disturbing the dam and was changed once a week thereafter. Animal experiments were approved by the Animal Care and Use Committee of the Jichi Medical University.

### 2.2 Genotyping

Mouse tails were collected, and genomic DNA was isolated by proteinase K digestion at 56°C overnight. The primer sequences used were as follows: *Neo* primers (sense) 5’-GTCCGGTGCCCTGAATGAACTGCAA-3’ and (antisense) 3’- ATTCGCCGCCAAGCTCTTCAG-5’. *V1bWT* primers: (sense) 5’- TCTGGCCACAGGAGGC AACCT-3’ and (antisense) 3’- ATCTCGTGGCAGATGAGGCCA-5’. The PCR protocol consisted of an initial denaturation step at 95°C for 5 min, followed by 30 cycles of denaturation at 94°C for 30 s and annealing/extension at 60°C for 30 s. PCR products were analyzed by agarose gel electrophoresis.

### 2.3 Behavioral test

#### 2.3.1 Free movement in open-field

We evaluated the locomotor and exploratory activities of male and virgin females at 8–12 weeks of age in a new PC cage (18 × 26 × 12 cm) with bedding. The behavior was recorded using a video camera (Victor, model GZ-MG77S) at 30 fps and 640 × 480 pixels/image without human interference for 15 mins. The test was performed between 9:00 and 12:00.

#### 2.3.2 Pup retrieval test

Maternal motivation to retrieve pups in a new environment was examined according to a previous report of the conventional condition with slight modifications^29^. On lactation day 2 (LD2), the pups were separated from their mother 60 mins prior to the test and kept on a thermopad (36°C) to maintain body temperature. Before testing, four pups were randomly selected and placed at the corners of the new cage (18 × 26 × 12 cm), which was covered by a handful of home cage bedding. The dam was placed in the center of the cage, and pup retrieval behavior was observed for 15 min. The number of retrieved pups was counted every minute. The tests were recorded using a digital video camera, and the files were used for deep learning experiments. After the test, the dams were brought back into their home cages with all the pups. The test was repeated on LD4 (n = 12 per genotype).

### 2.4 Machine learning models

#### 2.4.1 Preparation of images for training and evaluation of a deep learning network

RGB color images of 640 × 480 pixels were prepared from video files at a rate of one image per second using ffmpeg program^39^. To train and develop the first deep learning model, model 1, a total of 3044 images contained 740 images of dams and babies on clean beddings, 671 images of dams and babies in their home cages, and 1633 images of dams only in a cage used for the pup retrieval test. The ages of babies ranged from lactation day 1 to day 10 in both the WT and V1bKO groups. For the preparation of the second model (model 2), images of home cages were replaced with 800 images of pup retrieval tests, and a total of 3141 images were used. Dams and pups were defined as the objects in a training image and were boxed using LabelImg software (Tzutalin, https://github.com/tzutalin/labelImg). The total numbers of labeled objects were 5896 and 7062 for models 1 and 2, respectively. All images were randomly partitioned into training plus validation (89%) and test (9%) sets.

#### 2.4.2 Transfer learning and improvements of model performance

To develop a model for the detection of a mother and babies through computer vision, we used transfer learning based on a model of faster region-based convolutional neural networks (R-CNNs), which were pretrained on Microsoft Common Objects in Context (COCO) datasets and were included in the TensorFlow Object Detection API ^40–43^. For the region proposal network in Faster R-CNNs, ResNet101 was used ^40, 44^. The custom-built computer server worked with an Intel Core-i7-7800X 6 Core CPU and 32 GB RAM. For parallel calculations, a GeForce GTX1080Ti graphic card was used with CUDA 10.0 and 10.1 software for TensorFlow-gpu version 1.5 and TensorFlow version 2.3, respectively, in Ubuntu 18.04 operation system. A deep learning model was developed using the Miniconda package management system. Mouse images with labels were converted to TFrecords and used for training steps, according to the instructions of the TensorFlow Object detection API. The number of trains and evaluation steps were set to 50000 and 2000, respectively. Results from the training and evaluation processes were visualized in TensorBoard ^43^.

The center of the detection box was considered to be the XY coordinates of the corresponding object. Our Python code stores the distance matrix between each pair of two objects as a numpy array. The summation of distances between objects was calculated using the pdist function in the SciPy scientific computing program. Parameters, such as the number of boxes, total distance, and XY coordinates, were obtained as output, and were further analyzed using the R program. Our original code in R accepts three files for image name, number of boxes in an image, and total distance among objects. The number of boxes and total distance were plotted as Y-axis values and the number of the corresponding images as the X axis. Images were extracted one image per second from video data, thus the XY plot shows the time series of the relationship between mother and babies during the pup retrieval test. Distances of mother movement were accumulated, and compared between WT and V1bKO mice.

### 2.5 c-Fos-Immunohistochemistry

#### 2.5.1 Sample collection and tissue sectioning

After the pup retrieval test, the dams were kept alone in their home cages for 4 hrs, and then the pups were placed back into the same cage with the dam for another 6 hrs before being sacrificed. Lactating female mice were anesthetized by intraperitoneal injection of pentobarbital and perfused with ice-cold phosphate-buffered saline (PBS) with 4% paraformaldehyde (PFA) in PBS. The brains were collected and kept in 4% PFA overnight and then immersed in 30% sucrose at 4°C for 3 days. Brain sections with MPOA were collected, and 20-μm-thick coronal tissue sections were obtained using a cryostat (CM3050s, Leica, Nussloch, Germany) at −19°C.

#### 2.5.2 Immunohistochemistry analysis

The tissue sections were washed three times with 0.05% polysorbate 20 (Tween 20) in M PBS (PBS-T) and incubated in rabbit anti-*c-fos* primary antibody for 3 days at 4°C. After washing three times with PBS-T, the tissues were stained with a secondary antibody for 30 min and washed with PBS-T for 5 min. Sections were reacted with avidin-biotic peroxidase complex kit (Vectastain Elite Kit; Vector Laboratories) for another 30 min and washed in PBS-T for 5 min before staining with 3,3’-diaminobenzidine (DAB). After mounting the coverslips with Canada balsam mounting medium, *c-fos* expression was observed by light microscopy (AX80, Olympus,Tokyo, Japan), and quantitative results were analyzed using ImageJ.

### 2.6 In situ hybridization

WT brain tissues from virgin females and lactation day 2 mothers (n = 5) were immediately frozen in dry ice, and sectioned into 10-μm-thick slices using a cryostat for fluorescence *in situ* hybridization with ACDBio RNAscope^®^ multiplex fluorescence assay. The MPOA in the mouse atlas was located at bregma, ML ± 0.25 mm, DV - 5.00 mm^45^. The brain slices were fixed with 4% PFA at 4°C for 15 min and dehydrated with 50%, 70%, and 100% ethanol (2x) at room temperature for 5 min each; a 10-µm brain section was marked with a hydrophobic barrier pen. We used RNAscope^®^ protease IV (ACD bio, #322340) for pretreatment and RNAscope^®^ fluorescent multiplex detection reagents (ACD bio, #320851) to amplify the signal after probe hybridization. The positive target probes were a combination of Mm-Gal (#400961), Mm-Avpr1b-C2 (#480141-C2), and Mm-Fos-C3 (#316921-C3). DAPI staining was performed at the last step; a coverslip was mounted with prolong gold antifade antigens (Thermo Fisher Scientific, Eugene, OR USA) and then stored in a dark place at 4°C overnight. Signals from two slices were examined in each dam using a confocal laser microscope (FV1000; Olympus, Tokyo, Japan).

### 2.7 Statistical analysis

Data are expressed as the mean ± standard error of the mean (SEM). Statistical tests were performed using R software. The differences in pup retrieval numbers were analyzed using a two-way ANOVA. Two-tailed Student’s t-test or Wilcoxon test were used for single comparisons between the two groups. Statistical significance was set at *p* < 0.05.

## 3. Results

### 3.1 Deep learning analysis of freely moving mice

Before examining maternal behavior, we evaluated the free-moving behaviors of virgin female and male mice without infants in both WT and V1bKO groups. A rectangular animal cage was used as the open-field, because the mother-infant interactions were examined in the same cage in the subsequent experiments. To evaluate the mother-infant distances, TensorFlow Object Detection API^40–43^ was employed for training deep learning models; first (model 1) and second (model 2) models used 3044 and 3141 images, respectively, of dam and pup pairs from WT and V1bKO mice. Evaluation of the trained model 2 using 318 test images, which differed from those of the trained materials, resulted in a mAP score of 77% and total loss value of 0.45 in the TensorBoard program.

An adult female mouse was detected in a green box (Fig. 1a). The center of the detection box was traced as the position of the mouse (Fig. 1a and 1b). The total moving distances of female V1bKO mice in the open-field environment were significantly reduced, compared to those of WT mice. Figure 1c and 1d show individual and mean accumulated total distances, respectively. Time (F_1,18972_ = 65214.1, *p* < 0.001) and genotype (F_1,18972_ = 13272.9, *p* < 0.001) were significant determinants of the accumulated distance (Figure 1d). Two-dimensional views of the tracing of moving mice showed that V1bKO mice were less likely to enter the centerfield (Fig. 1e). Accumulated counts of centerfield entrance were significantly low in V1bKO mice, compared with those of WT mice (Fig. 1f, *p* = 0.0344). In Figure 1g, red points indicate that V1bKO females entered the centerfield less frequently than WT, thereby indicating more anxiety-like behavior in V1bKO (*p* = 0.0042). Our trained model detected both female WT and V1bKO mice equally well; the detection failure rates were 0.14% and 0.21% (*p* = 0.06, n = 10 for WT and V1bKO mice).

**Fig. 1.**
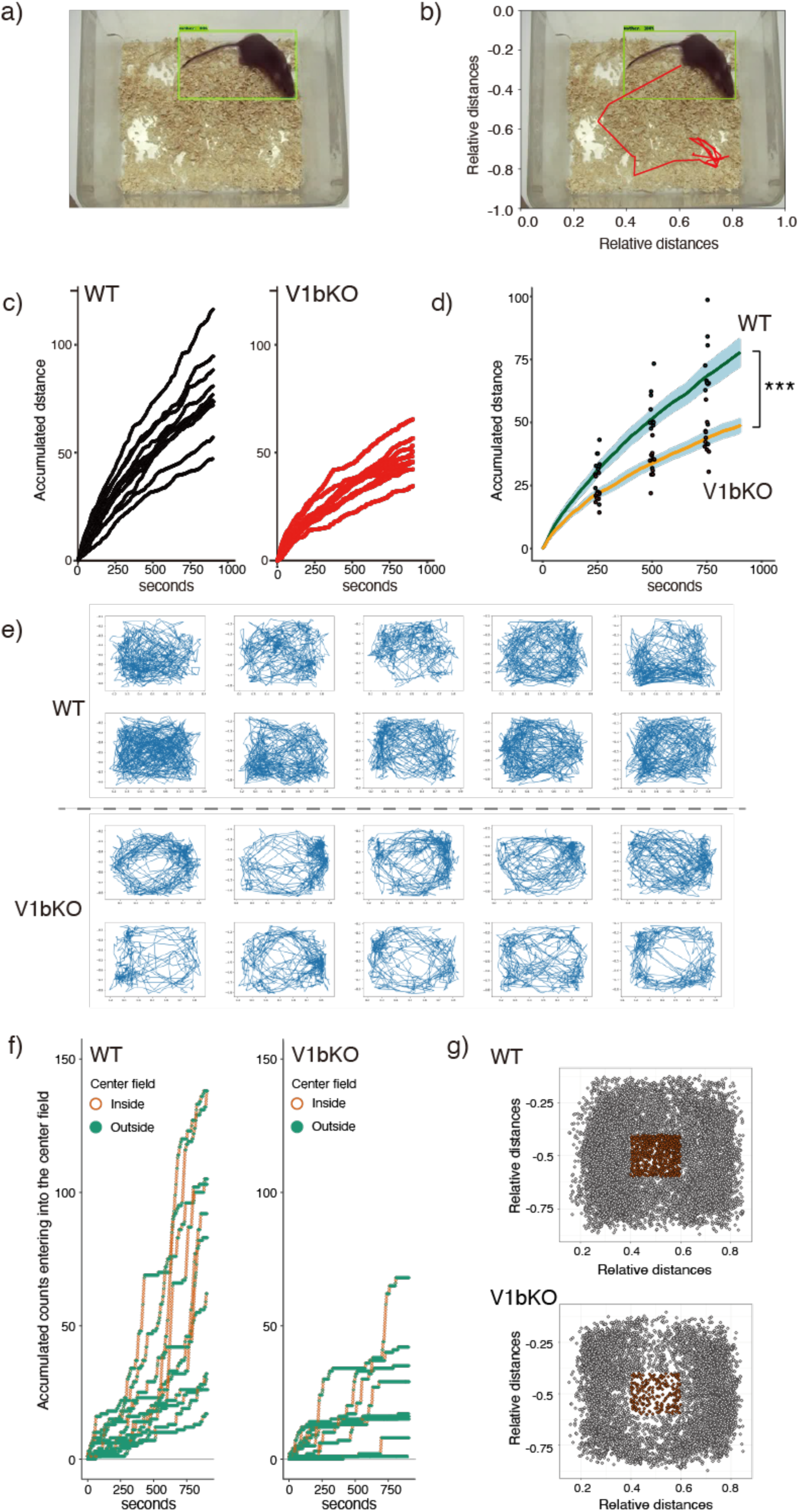
Object detection and tracking using a deep neural network revealed anxiety-like behaviors in V1bKO female mice in the open field. (a) Images from video recordings of free moving mice were obtained at one image per second. Mice in the images were located with a bounding box (green). (b) The XY coordinates of the center of the detected box were traced to calculate total distance. (c) Total distance was accumulated and plotted for individual mouse in WT (black) and V1bKO (red) groups, respectively. (d) Mean ± SEM of accumulated total distances in WT (n=10) and V1bKO (n=10). Data points of individual mice were plotted at the 250^th^, 500^th^, and 750^th^ images to show their distribution. ***, *p* < 0.001. (e) Images show two-dimensional traces of the open-field test. V1bKO mice entered in the centerfield less frequently, compared to WT mice. (f) The width and height of the entire open-field environment were set as one and the centerfield was defined between 0.4 and 0.6 of the relative distances in both x and y axis. Detection in the centerfield led to one point, and points were accumulated (orange) in sequential images in the open-field test. Detection of the mouse outside the centerfield led to zero points (green). (g) Red points indicate the positions of mice, which were detected in the centerfield. The results are from all mice examined in this experiment.

As observed in the female V1bKO mice, male V1bKO mice explored less distance in the open-field, compared with WT mice (Fig. 2a and 2b for individual and mean accumulated distances, respectively). In Fig. 2b, time (F_1,18113_ = 98045.5, *p* < 0.001) and genotype (F_1,18113_ = 3895.3, *p* < 0.001) were significant determinants of the accumulated distance. However, in contrast to the female V1bKO mice, the difference between male V1bKO and WT mice in the two-dimensional tracing was not apparent in the open-field (Fig. 2c). The accumulated mouse detection counts in the centerfields were similar between male V1b and WT mice (Fig. 2d, *p* = 0.716). Also, the points where male mice were detected in the center area were similar between the two groups (*p* = 0.9627).

**Fig. 2.**
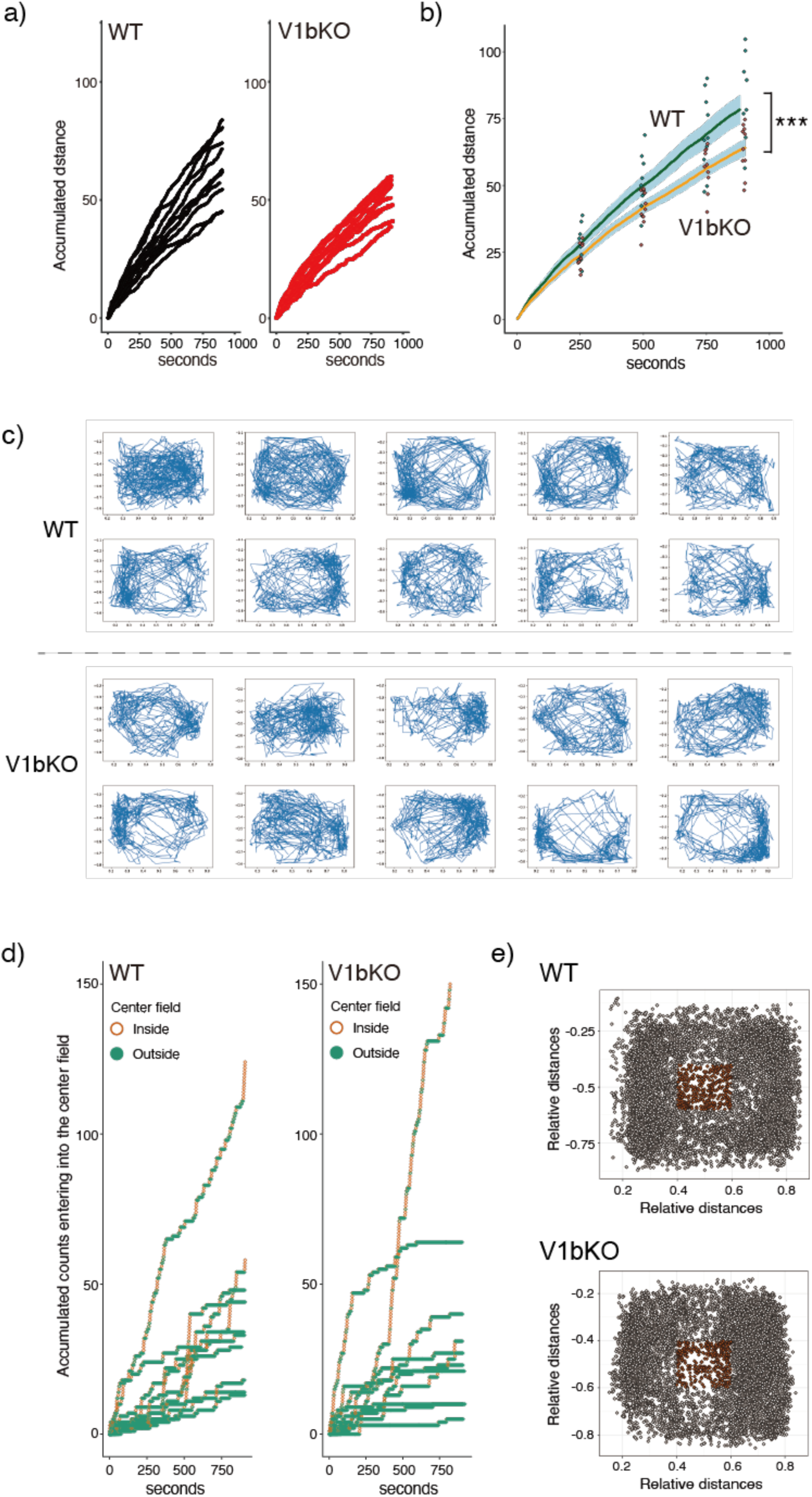
Male V1bKO mice explored less distance, compared to WT mice, but both mice entered the centerfield at similar frequencies. (a) Total distances traveled by individual mice were plotted against the number of sequential images, obtained at one image per second. (b) Mean ± SEM of accumulated total distances moved in WT (n = 10) and V1bKO (n = 10). Data points of individual mice were plotted on the 250^th^, 500^th^, and 750^th^ images to show their distributions. ***, *p* < 0.001. (c) Images show two-dimensional traces of traveling mice in the open-field test. (d) Entrance incidents into the centerfield were counted and accumulated counts were plotted using orange. Staying outside the centerfield was plotted in green. (**e**) Points, where mice were detected in the open-field tests, were plotted using red if the mice stayed in the centerfield and using gray if the mice were outside the centerfield.

To ensure consistency in our object detection protocol, we compared the results from repeated detections and those from different versions of TensorFlow programs (version 1.15 vs. 2.4). Repeated detection resulted in the same mouse positions and tracing lines in both TensorFlow versions (Supplemental Fig. 1a and 1b). When two versions of the TensorFlow programs were trained separately using the same set of images and labels, a similar tendency was observed (Supplemental Fig. 1c). However, the detected positions or lines between the objects were not the same (Supplemental Fig. 1c). Therefore, we used the same TensorFlow version (version 1.15) to compare the WT and V1bKO in this study, unless otherwise stated.

**Supplemental Fig. 1.**
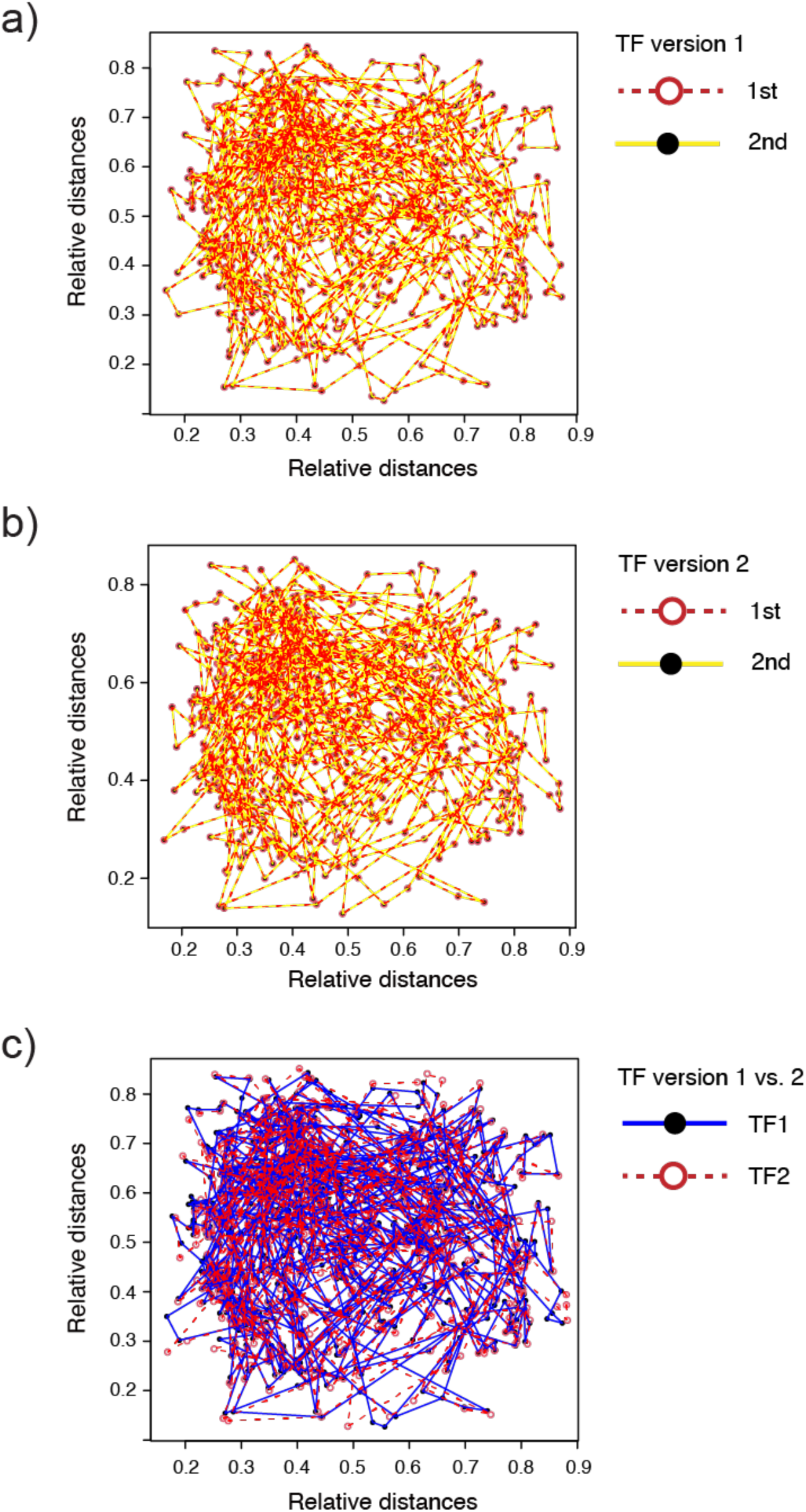
Repeated detection resulted in the consistent localization of the freely moving mouse. A set of 910 images was repeatedly detected by using deep learning models developed by the TensorFlow program version 1.15 (SFig. 1a) and 2.4 (SFig. 1b). The locations of the centers of the detection boxes are indicated to be using dots (first detection) and red circles (second detections), respectively. Tracings of ambulation in the first and second detections are indicated by red dashed lines and yellow lines, respectively. (SFig. 1c) Detections by different TensorFlow versions were compared between versions 1.15 (black dots and blue lines) and 2.4 (red circles and red dashed lines).

### 3.2 Pup retrieval test: manual measurement

To examine the role of V1b in pup-directed maternal behavior during the early lactation period, pup retrieval test was performed on lactation days 2 and 4. In pup retrieval in rodents, dams seek their pups, gently pick them up by mouth, and bring them to the nest ^46^. After 60 min of separation from the mother, four pups were placed at each corner of a new cage. After introducing the mother at the center of the cage, the retrieval behavior was counted every minute for 15 mins. The entire test was recorded using a video camera. Manual counting of pup retrieval showed that the V1bKO mother collected her pups faster than the WT mother (Fig. 3). In Fig. 3a and 3b, both mouse genotype (*p* < 0.05) and time (*p* < 0.001) were significant determinants of the number of retrieved pups in the two-way ANOVA analysis.

**Fig. 3.**
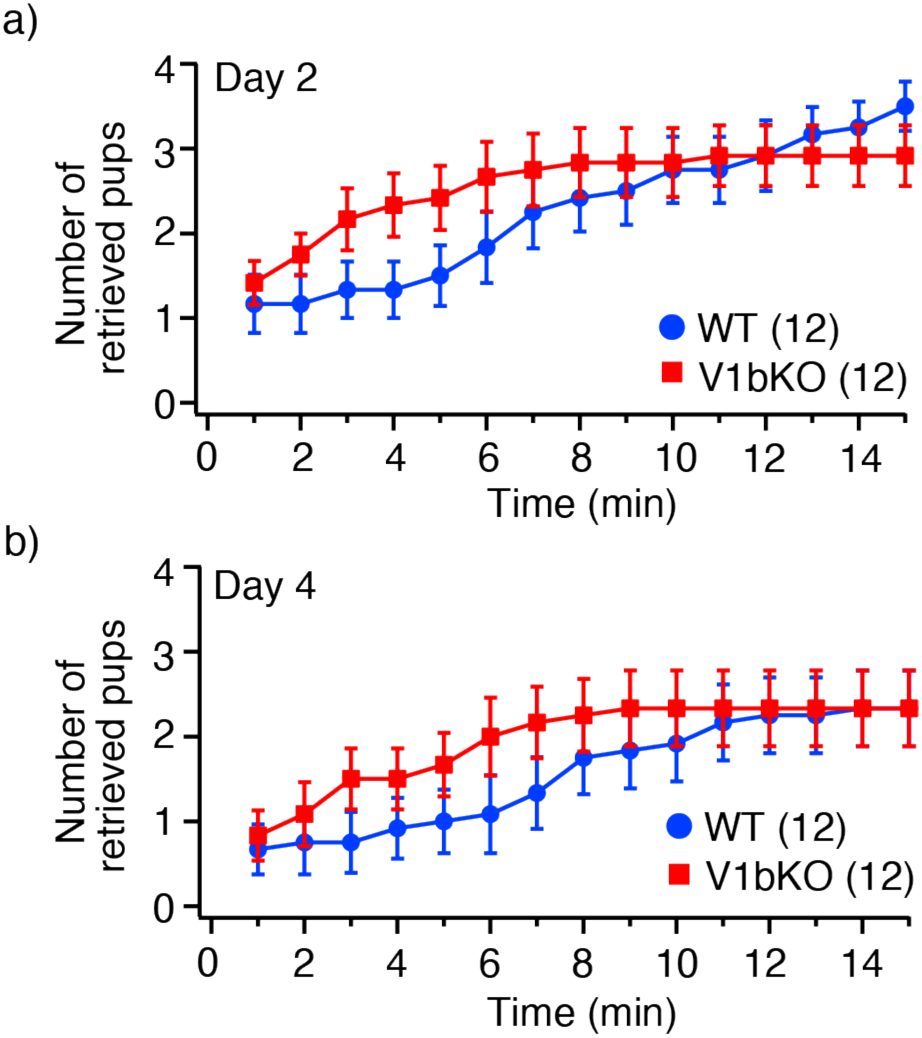
V1bKO mother retrieved their pups faster than WT mothers in the manual measurement of the pup retrieval test. Prior to the transfer of the mothers to the novel test cage environment, four pups were separated from their mothers for 60 mins and placed at separate corners of the cage. The mother was placed at the center of the field and the mother’s retrieval behavior was counted every minute for 15 mins on their lactation day 2 (a) and day 4 (b). Data are presented as mean ± SEM (n = 12 per group).

### 3.3 Deep learning analysis of dam-pups interaction

The detection accuracy of our deep learning model was 99% in both WT (1825/1829) and V1bKO (1832/1842) mice (Supplemental Fig. 2).

**Supplemental Fig. 2.**
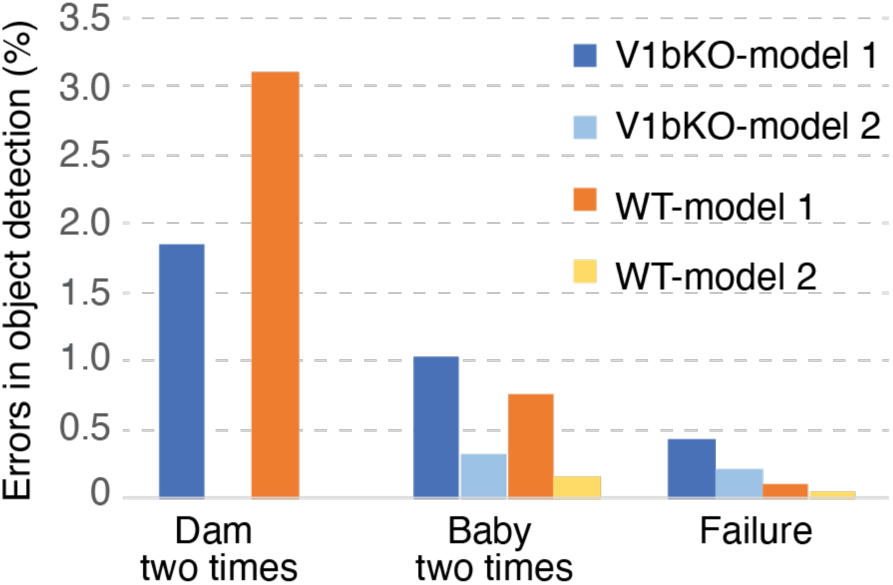
Error rates of object detection by the first and second models. Same sets of 1829 and 1842 images of WT and V1bKO mother-baby pairs, respectively, were detected by the first and second deep learning models to evaluate detection accuracy. The results of the bounding box and the detected objects were manually evaluated, and the detection errors were indicated. To avoid the detection of a mother twice in the second model, the mother’s XY coordinate of the highest probability was selected.

In the object detection, the height and width of the two-dimensional image were set to 1, to ensure that the positions of the objects were expressed as (0 < x < 1, 0 < y < 1). At the beginning of the retrieval test, pups at each corner and the dam made five objects (Fig. 4a), and the sum of the distances between each pair of objects tended to decrease based on the progress if the dam collected her pups effectively (Fig 4b and 4c). On lactation day 2, the accumulated total distance (Fig. 4d and 4e) and the number of objects (Fig. 4f and 4g) during the test were similar between the two mouse groups (*p* = 0.471, and *p* = 0.665 for total distance and box numbers, respectively). However, on lactation day 4, the accumulated total distance between the objects was reduced in V1bKO mice compared to WT mice (Fig. 4h and 4i. *p* = 0.0172 and *p* < 0.001 for total distance and box numbers, respectively). Correspondingly, the number of detected objects decreased more quickly in V1bKO mice than in WT mice on day 4 (Fig. 4j). Moreover, when object numbers were accumulated during the test, the data from V1bKO mice included significantly fewer objects than the data from WT mice on day 4 (Fig. 4k). Therefore, our deep learning analysis of the pup retrieval test identified the achievement of closer interactions with pups by the V1bKO dam, compared to the WT dam on lactation day 4.

**Fig. 4.**
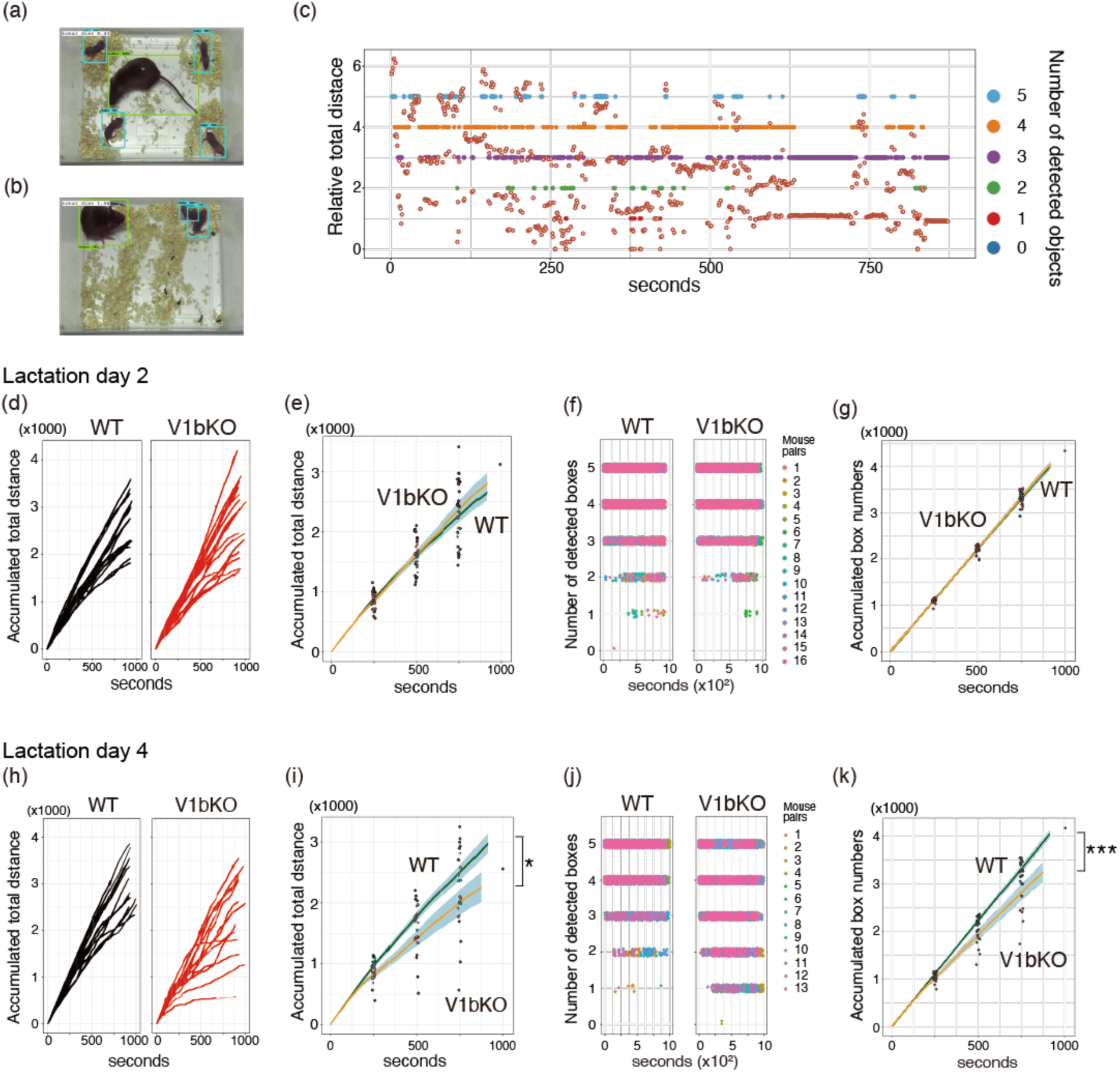
Deep learning analysis of pup retrieval test revealed shorter distances between V1bKO dam and pups at lactation day 4. The distances between all pairs of objects in an image were accumulated to show the approximate distribution of dam and pups on lactation days 2 and 4. For example, the total distances were 8.43 and 1.56 in the first (a) and 658^th^ (b) images, respectively. (c) Time course of total distances (red circles and left Y-axis) and number of detected boxes (from 1 to 5 and right Y-axis) were plotted against the time (one image per second). The total distances between pairs of objects were accumulated during the retrieval test and individual (d and h) and mean accumulated distances and SEM (e and i) were plotted. The numbers of recognized objects in an image (f and j) or mean ± SEM values of the accumulated object numbers (g and k) were plotted against time. In panel (j), object numbers come to one, which means effective retrieval of four pups by a V1bKO dam. Data are derived from 16 and 13 dam-pup pairs in both mouse groups on lactation day 2 and 4, respectively. *, *p* < 0.05. ***, *p* < 0.001.

In the pup retrieval test, traces of the dam’s movements in a two-dimensional view revealed a reduction in the movements of V1bKO dams on both days 2 and 4 (Fig. 5a and 5b). The accumulated total moving distances of individual mice (Fig. 5c and 5d) and mean values (Fig. 5e and 5f) confirmed reduced moving distances in V1bKO dams. In both mouse types, the accumulated total distances were reduced on lactation day 4, compared with those on day 2. Histograms of moved distances in one frame show that V1bKO moved in short distances more frequently than WT mice on both lactation days (Fig. 5g and 5h). These results indicate that V1bKO dam effectively retrieved their pups in shorter total movements compared to WT dam.

**Fig. 5.**
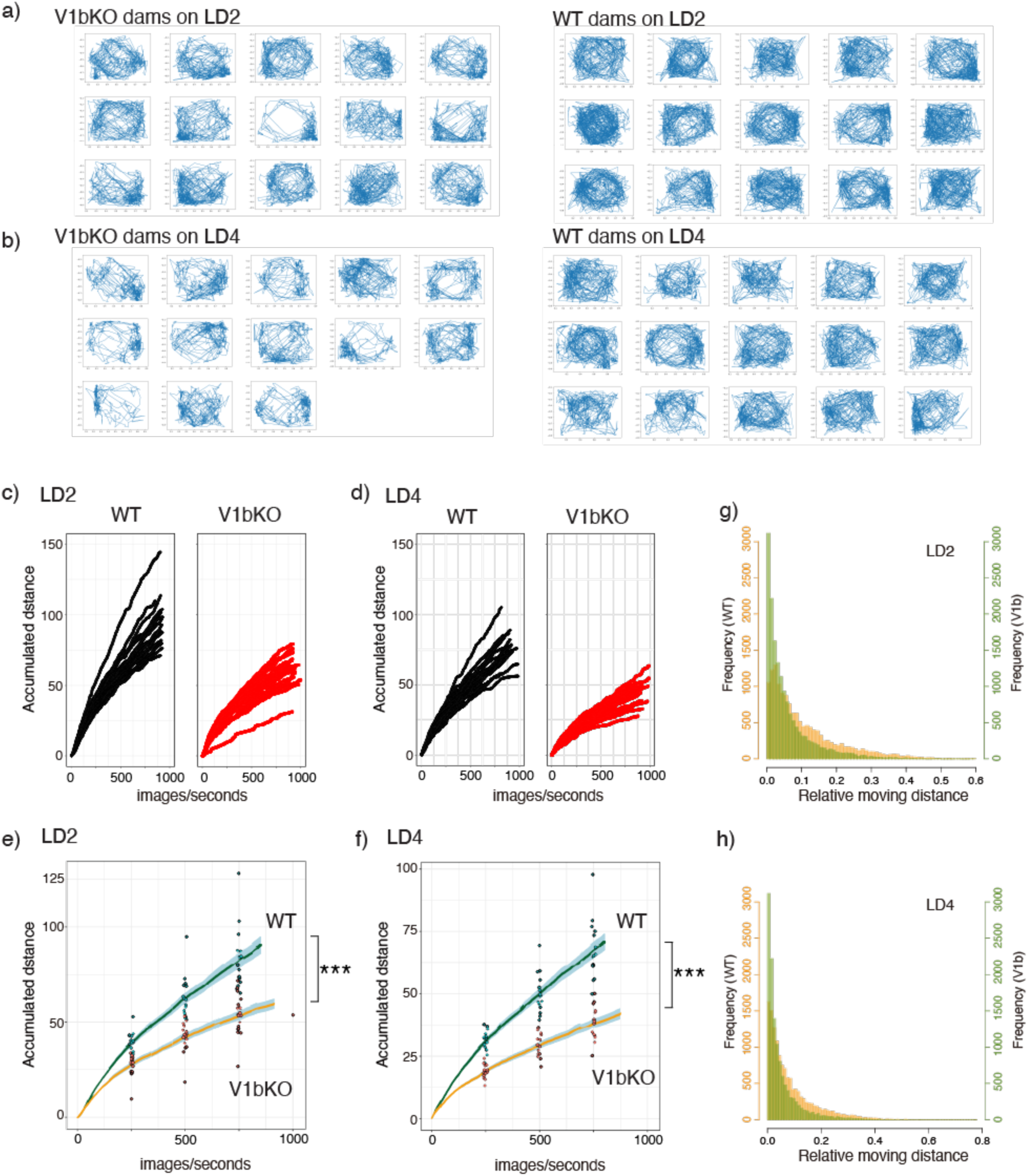
V1bKO dams moved less distances per time during the pup retrieval test. Movements of dams during the pup retrieval test were obtained as two-dimensional tracings on lactation day 2 (a) and lactation day 4 (b). Accumulated total distances of individual dams’ movement were plotted against time at lactation day 2 (c) and day 4 (d). (e) and (f) show the mean ± SEM of the accumulated total distances from the data in (c) and (d), respectively. ***, *p* < 0.001. Histograms show the frequencies of moved distances in a flame/sec in WT (orange) and V1bKO (green) dams on lactation day 2 (g) and day 4 (h).

### 3.4 Histological examination of the MPOA

Because functional deletion of V1b gene altered pup retrieval behavior, we further examined neuronal activities and V1b receptor mRNA expression in the MPOA of postpartum female mice.

To examine *c-fos* expression as a neuronal activity in MPOA ^48–50^, we initially separated pups from their dam for 4 hrs; then, the pups were reunited with the dam as stimuli for 6 hrs. The results showed that the number of *c-fos* positive cells was significantly lower in V1bKO dams than in WT dams (Fig. 6a and 6b, Wilcoxon, \**p* = 0.0087, n = 6 and 5 for V1bKO and WT, respectively). In contrast, the *c-fos* positive nuclei at cingulate cortex area 2 (Cg2, Fig. 6j) were not different between WT and V1bKO mice (Fig. 6i, Wilcoxon, \**p* = 0.12). In galanin-positive and -negative neurons in the MPOA of WT virgin female mice and dams on lactation day 2, the number of V1b transcript-positive neurons was similar independent of the status of the mother’s pregnancy (Fig. 6e, n = 5). The V1b transcript was detected in both galanin-positive and galanin-negative neurons (Fig. 6f and 6g, n = 5 in each group). c-Fos positive signals were detected in the galanin-V1b double positive cells and galanin-negative and V1b-positive cells (Fig. 6h). The total numbers of V1b expressing cells were not different between virgin female and lactating mothers (371 and 428 cells for virgin females and lactating dams, respectively; Wilcoxon test, *p* > 0.05).

**Fig. 6.**
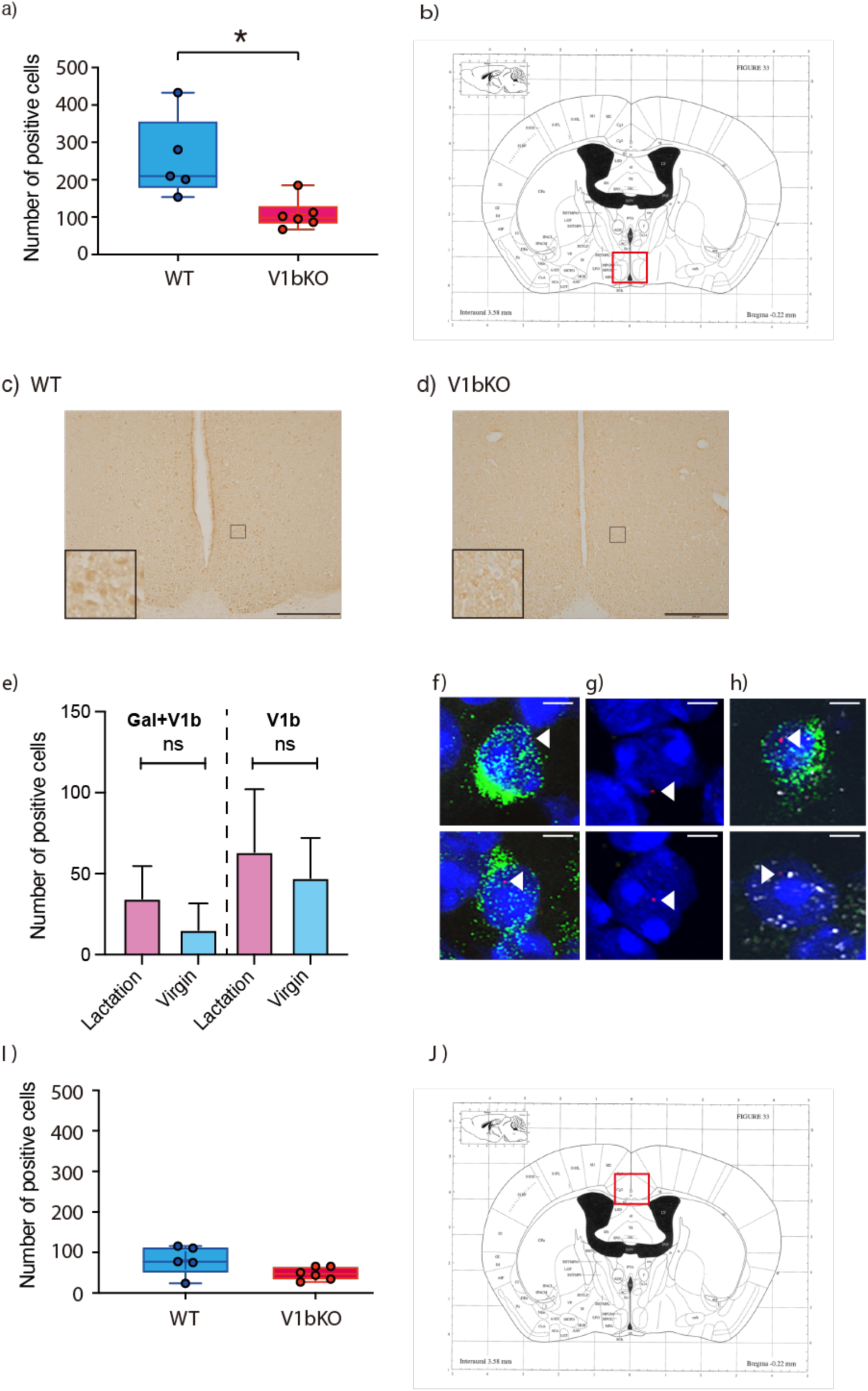
*c-Fos* and V1b expression in the MPOA. (a) After stimulation of dams by pups, the *c-fos* positive nuclei in the MPOA were examined in WT and V1bKO mice. *, Wilcoxon test, *p* = 0.0087 (n = 5–6). (b) A red square in the brain map shows an area, where gene expression was examined. The brain map was from cited reference^47^. (c) and (d) In immunohistochemistry of MPOA, *c-fos* signals after pup stimulation were weaker in V1bKO dams compared to WT dams. Insets show large magnification of the cells. Scale bars indicate 200 µm. (e) *In situ* hybridization identified expression of V1b mRNA in galanin-positive and -negative cells in the MPOA of WT virgin mice and mother mice on postpartum day 2. Wilcoxon test, ns, *p* > 0.05. Expression of V1b (red) was indicated with white arrowheads in galanin (green)-positive (f) and -negative (g) neurons. (h) After pup stimuli, *c-fos* (white) and V1b (red) double-positive cells with (upper panel) or without (lower panel) galanin expression (green) were identified in the MPOA of WT mother. Scale bar indicates 5 µm. (i) The number of *c-fos* positive nuclei was not different in cingulate cortex area 2 (j) in the two mouse groups.

## 4. Discussion

In this study, deep learning and computer vision delineated the role of V1b in mother-pup interaction by closely analyzing the pup retrieval tests. V1b receptor was necessary to actively explore the open-field in both male and female mice; lack of the V1b gene resulted in shorter moving distances compared to WT. Regarding the frequency of entering the centerfield, sex differences were detected. The V1b was necessary only for female mice, not male mice, to enter the centerfield. This phenotypic tendency of virgin V1bKO females was also maintained in lactating V1bKO mothers. Unexpectedly, the V1bKO dams with less movement retrieved mice more efficiently than the WT dams. The deep learning model was effective in evaluating distances between mother and pups and between pups altogether.

Furthermore, the number of detected objects was a useful measure for the efficiency of collection, because after the pups were collected, the dam-covered pups and the dam were counted as one object. Our findings further suggest that the V1b receptor, which is expressed in the MPOA, especially in parts of galanin neurons, may play a role in modulating the efficiency of maternal care in pup retrieval.

In an open-field test, virgin female V1bKO mice moved less distance and entered the center of the cage less frequently than WT mice. Behaviors of avoiding an entry to the centerfield or of walking close to the wall indicated high anxiety-like emotional states ^50–52^. A previous study on male V1bKO and WT mice reported no change in the open-field test in the frequency of total beam breaks ^53^. Several experimental conditions are different between the previous and current studies. The open-field test in the previous study was performed during dark phase, but we examined during the daytime according to the report by Bayerl et. al^29^. In addition, the floor of the field was covered with a small amount of wood shavings for bedding. Therefore, the condition of the bedding materials might impact mouse behavior ^54, 55^.

The effect of V1b deletion on pup retrieval is in good agreement with a previous study, which reported that blockage of the V1b receptor by local administration of the antagonist into the MPOA, but not into the lateral ventricle, led to faster pup retrieval^29^. AVP has been shown to have a profound effect on maternal behavior through the V1a and V1b receptor in the MPOA^56^. Oligonucleotide-mediated knockdown of V1a reduced the number of retrieved pups^18^. In contrast, the acute effect of the V1a antagonist on BNST did not change time course of pup retrieval^57^. Therefore, AVP in the MPOA is likely to have an opposite effect on pup retrieval through the V1a and V1b subtypes. Both of these V1 receptors couple with Gq protein and cause Ca^2+^ signaling in the cell ^28^. However, coupling with β-arrestin, receptor localization in the cell, and recycling behavior after stimulation by AVP differ remarkably^58–61^. It is of interest to understand how sibling receptors exhibit opposite phenotypes in maternal behavior, especially in pup retrieval.

A part of the V1b receptors in the MPOA was expressed in galanin-positive neurons, and those neurons were activated and *c-fos*-positive in the lactating dam. *c-Fos* expression is significantly increased in the MPOA of dams when dams are stimulated by pups^49, 62^. We noted that overall V1b receptor mRNA levels did not change between lactating dams and virgin female mice. Similarly, V1b receptor mRNA and protein expression levels did not change in the MPOA or BNST of lactating rats ^29^. In V1bKO mice, the number of *c-fos*-positive neurons in the MPOA was decreased. Because not all *c-fos*-positive cells express the V1b receptor, reduction of *c-fos*-positive cells can be due to direct as well as indirect effects of V1b deletion. A subset of galanin neurons are active during the maternal period, and genetic deletion of these neurons significantly reduced pup retrieval behaviors^10^. Elucidation of possible relationship between galanin neuron and V1b receptor function is of interest and needs further study. The V1b receptor might negatively regulate pup retrieval, and deletion of the V1b resulted in an enhanced retrieval in a new environment. A recent comprehensive work on V1b and OT receptor double KO mice supports a part of this idea. After a stressed condition, reduced pup care by OT receptor KO mother was recovered by further deletion of V1b gene ^63^. Therefore, functional interaction between V1b and other receptors needs to be considered.

Deep-learning-based behavioral analysis enables precise evaluation of gene and drug effect on experimental and wild animals ^64^. In the analysis of mothers and pups by deep neural network and computer vision, the size and color are useful features for the separate detection of objects. However, pups continue to change in appearance after birth, which makes it challenging to continuously detect pup locations. We found that training pup images from parturition to ten days of age every day effectively establish a deep learning model, which can be applied on both days 2 and 4 equally. After obtaining the XY coordinates of the mother and pups, the efficacy of pup retrieval was evaluated by two parameters, the accumulated distance between each object and the number of objects. The accumulated distance between each object indicates how the objects were spread in the test field. The number of objects at the beginning is five: four pups and mother; this number changes to one, if the mother collects all pups and covers them under her. A plot of the time series of these two parameters during pup retrieval showed a correlation between them, as shown in Fig. 4c. Interestingly, the V1bKO dams moved less distance, and entered the center of the field less often than WT dam. However, retrieval of the pups was faster in V1bKO dam than in WT dam. During the explorations of new environment, the WT mice encountered pups several times; however, retrieval behaviors were not initiated. Less moved distance in V1bKO dam indicates that a balance between exploration and goal-directed, pup retrieval behavior might be different between V1b and WT dams. Following these results, the next question to address isto investigate how these two groups of mice exhibit maternal care in their home cages. It is challenging to perform lasting recording and deep learning analysis of large records. However, the results of the current study and recent reports hold promise for further extending this research between mothers and infants ^38^. Relying on deep learning to make a judgment model is robust, as we have showed here. In addition, the reproducibility of the results is clearly shown as another advantage of deep learning and computer vision.

## 5. Conclusion

In summary, the V1b receptor plays a crucial role in both exploring new environments and in the retrieval of pups by lactating mouse mothers. This knowledge from deep learning analysis is useful to help understand the efforts of parents when fostering their infants during the early phase of the postpartum period.

## Acknowledgements

This work was supported, in part, by Grants-in-Aid for Scientific Research from the Ministry of Education (T.K.), The Science Research Promotion Fund (T.K.), JKA through its promotion funds from KEIRIN RACE (T.K.), and Jichi University Young Investigator Award (T.K.).

## Author contributions statement

S.C. and T.K. conceived the project and wrote the manuscript. S.C., F., and A.M. performed the experiments. S.C. and T.K. designed the experiments. All authors analyzed the data. T.K. coordinated and directed the project. All authors reviewed the manuscript.

## Data and materials availability

The datasets generated during and/or analyzed during this study are available from the corresponding author on reasonable request. All unique/stable reagents generated in this study are available from the Lead Contact upon the completion of a Materials Transfer Agreement.

## Code availability

The code used during this study are available from the corresponding author on request. Image data and its analysis code will be deposited to Scientific Data.

